# Anaplerotic flux into the Calvin-Benson cycle. Hydrogen isotope evidence for *in vivo* occurrence in C_3_ metabolism

**DOI:** 10.1101/2021.07.30.454453

**Authors:** Thomas Wieloch, Angela Augusti, Jürgen Schleucher

## Abstract

- As the central carbon uptake pathway in photosynthetic cells, the Calvin-Benson cycle is among the most important biochemical cycles for life on Earth. A carbon flux of anaplerotic origin (i.e., through the chloroplast-localised oxidative branch of the pentose phosphate pathway) into the Calvin-Benson cycle was proposed recently.
- Here, we measured intramolecular deuterium abundances in leaf starch of *Helianthus annuus* grown at varying ambient CO_2_ concentrations, *C*_a_. Additionally, we modelled deuterium fractionations expected for the anaplerotic pathway and compared modelled with measured fractionations.
- We report deuterium fractionation signals at H^1^ and H^2^ of starch glucose. Below a *C*_a_ change point, these signals increase with decreasing *C*_a_ consistent with modelled fractionations by anaplerotic flux. Under standard conditions (*C*_a_=450 ppm corresponding to intercellular CO_2_ concentrations, *C*_i_, of 328 ppm), we estimate negligible anaplerotic flux. At *C*_a_=180 ppm (*C*_i_=140 ppm), more than 10% of the glucose 6-phosphate entering the starch biosynthesis pathway is diverted into the anaplerotic pathway.
- In conclusion, we report evidence consistent with anaplerotic carbon flux into the Calvin-Benson cycle *in vivo*. We propose the flux may help to (i) maintain high levels of ribulose 1,5-bisphosphate under source-limited growth conditions to facilitate photorespiratory nitrogen assimilation required to build-up source strength and (ii) counteract oxidative stress.

## Introduction

In photosynthetic cells, carbon is taken up primarily by the Calvin-Benson cycle (CBC). Thus, the CBC is among the most important biochemical cycles for life on Earth. Sharkey and Weise (2016) proposed the CBC may be anaplerotically refilled by carbon injection from the oxidative branch of the chloroplast-localised pentose phosphate pathway (aka., glucose-6-phosphate shunt, orange pathway in Fig. 1). This pathway liberates CO_2_ from metabolism and anaplerotically recovered energy (NADPH) is insufficient for quantitative refixation. Should this flux occur *in vivo*, it would therefore affect plant carbon and energy balances, plant performance, ecosystem productivity, and biosphere-atmosphere CO_2_ exchange. This includes the CO_2_ fertilisation effect which has been identified as major unknown in our current Earth system understanding (IPCC, 2021) as well as future atmospheric CO_2_ concentrations and crop yields (*cf.* Long *et al.*, 2006).

**Figure 1.**
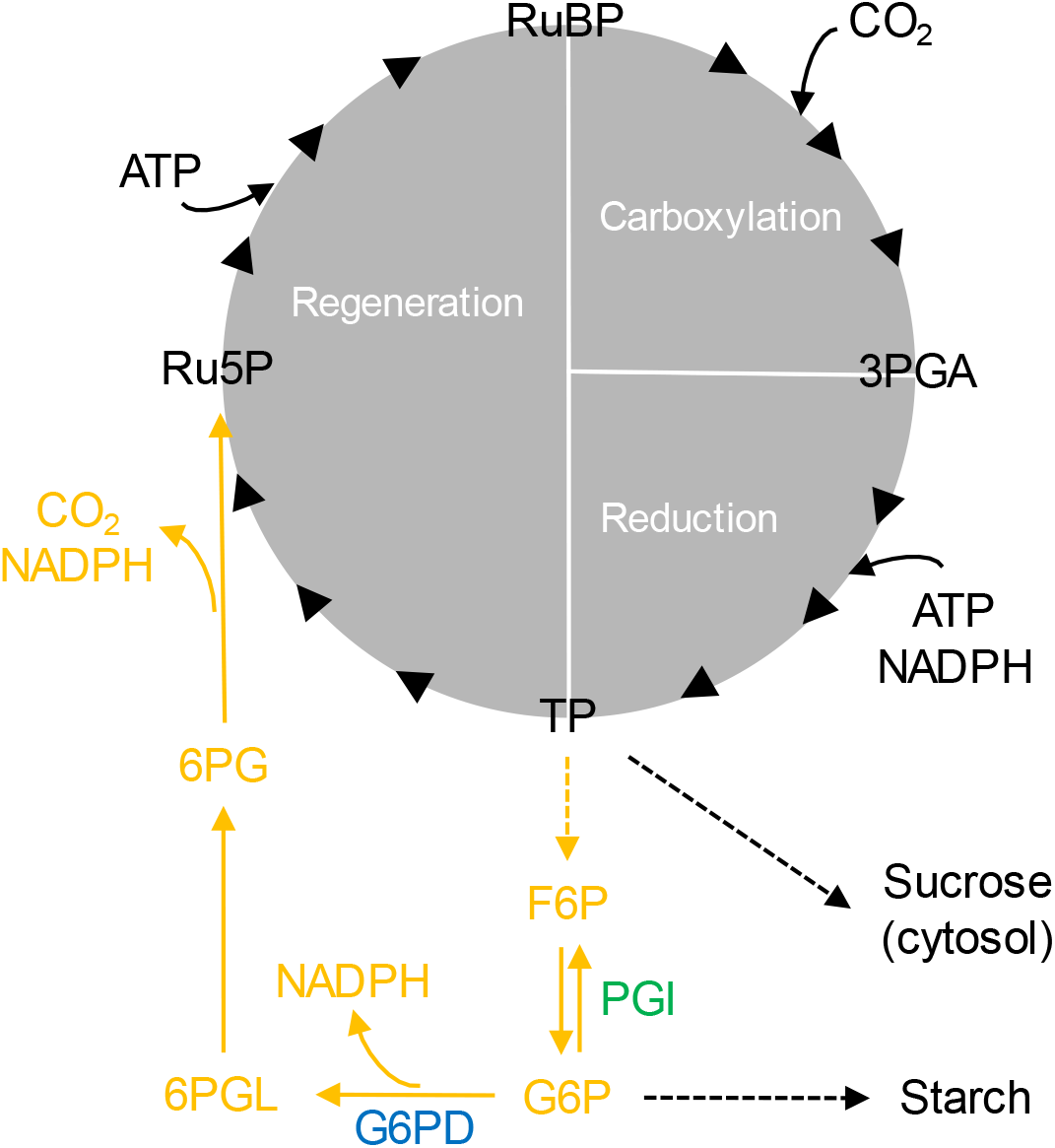
Oxidative branch of the chloroplast-localised pentose phosphate pathway carrying anaplerotic carbon flux into the Calvin-Benson cycle (in orange). Blue and green: Enzyme reactions that may introduce D fractionation signals at glucose H^1^ and H^2^, respectively. Dashed arrows: Intermediate reactions not shown. Enzymes: G6PD, glucose-6-phosphate dehydrogenase; PGI, phosphoglucose isomerase. Metabolites: 3PGA, 3-phosphoglycerate; 6PG, 6-phosphogluconate; 6PGL, 6-phosphogluconolactone; ATP, adenosine triphosphate; F6P, fructose 6-phosphate; G6P, glucose 6-phosphate; NADPH, nicotinamide adenine dinucleotide phosphate; Ru5P, ribulose 5-phosphate; RuBP, ribulose 1,5-bisphosphate; TP, triose phosphates (glyceraldehyde 3-phosphate, dihydroxyacetone phosphate). Modified figure from Wieloch *et al.* (2021).

Anaplerotic flux is believed to be controlled at phosphoglucose isomerase (PGI, EC 5.3.1.9; green in Fig. 1) which catalyses interconversions of fructose 6-phosphate (F6P) and G6P (Sharkey & Weise, 2016). In the light, the reaction is strongly displaced from equilibrium on the side of F6P keeping chloroplastic G6P concentrations low (Dietz, 1985; Gerhardt *et al.*, 1987; Kruckeberg *et al.*, 1989; Schleucher *et al.*, 1999) proposedly to restrict the anaplerotic flux (Sharkey & Weise, 2016). With decreasing PGI inhibitor concentrations (3-phosphoglycerate, and especially erythrose 4-phosphate), the PGI reaction shifts towards equilibrium (Dietz, 1985; Backhausen *et al.*, 1997). Concomitantly increasing G6P concentrations (Dietz, 1985) may cause anaplerotic flux increases by the following mechanism (Sharkey & Weise, 2016). In the light, the first enzyme of the pentose phosphate pathway, glucose-6-phosphate dehydrogenase (G6PD, EC 1.1.1.49; blue in Fig. 1), is inhibited by redox regulation via thioredoxin (Née *et al.*, 2009). However, inhibition can be reversed allosterically by increasing G6P concentrations (Cossar *et al.*, 1984; Preiser *et al.*, 2019).

While anaplerotic flux into the CBC seems biochemically feasible, so far, only three studies have reported flux-level evidence. First, Wieloch *et al.* (2018) analysed intramolecular ^13^C/^12^C ratios in glucose extracted from an annually resolved *Pinus nigra* tree-ring series (1961 to 1995). They reported ^13^C fractionation signals (i.e., systematic ^13^C/^12^C variation) at glucose C-1 and C-2. These signals respond to drought proposedly due to changes in anaplerotic flux by the following mechanism (Wieloch *et al.*, 2018). As isohydric species, *Pinus nigra* responds to drought by stomatal closure which impedes CO2 uptake causing low intercellular CO2 concentrations, *C*_i_ (Sade *et al.*, 2012). With decreasing *C*_i_, PGI inhibitor concentrations reportedly decrease, and the chloroplastic PGI reaction shifts towards equilibrium (Badger *et al.*, 1984; Dietz & Heber, 1984; Dietz, 1985). Concomitantly increasing G6P concentrations can be expected to cause increasing G6PD activity and anaplerotic flux (see last paragraph). Based on reported ^13^C isotope effects (Gilbert *et al.*, 2012), Wieloch *et al.* (2018) predicted ^13^C/^12^C increases at C-1 and C-2 at a ratio of 2.25 in response to shifts of the PGI reaction towards equilibrium and found a ratio of 2.74 (+1.35SE, −0.60SE) confirming their prediction. Interestingly, the reported ratio is somewhat higher than the predicted ratio yet not significantly. Still, the offset may be explained by the ^13^C effect of G6PD (*α*=1.0165; Hermes *et al.*, 1982) which can be expected to cause an additional ^13^C/^12^C increase at C-1 as anaplerotic flux increases.

Second, Wieloch *et al.* (2021) also analysed intramolecular deuterium (D) abundances across the *Pinus nigra* tree-ring series. They reported two closely related D fractionation signals at glucose H^1^ and H^2^ which respond to drought and atmospheric CO_2_ concentrations. Interestingly, the response requires the crossing of a change point. Wieloch *et al.* (2021) hypothesise that shifts of the chloroplastic PGI reaction towards equilibrium contribute to the signal at H^2^ while chloroplastic G6PD contributes to the H^1^ signal as anaplerotic flux changes.

Third, Xu *et al.* (2021) analysed ^13^C enrichment patterns of central metabolites in *Camelina sativa* leaves by ^13^C isotopically nonstationary metabolic flux analysis. They reported estimates for anaplerotic flux (4.6 μmol CO_2_ g^−1^ FW hr^−1^, ≈3% of Rubisco carboxylation) and day respiration (5.2 μmol CO_2_ g^−1^ FW hr^−1^). However, these estimates come from a physiologically unrealistic model and could not be confirmed by independent methods (Wieloch, 2021). In summary, Xu *et al.* (2021) reported evidence for anaplerotic flux under normal growth conditions while Wieloch *et al.* (2018, 2021) reported evidence for stress-induced upregulation.

To explain the ^13^C and D fractionation signals observed in tree-ring glucose of the gymnosperm *Pinus nigra*, Wieloch *et al.* (2021) proposed the following hypotheses. First, anaplerotic flux into the CBC is associated with D fractionation signals at H^1^ and H^2^ of chloroplastic G6P and its derivatives including leaf starch. Second, anaplerotic flux and associated D signals increase with decreasing *C*_i_. Third, increases occur below a *C*_i_ change point. In addition, these authors discussed whether anaplerotic flux is a general feature of C_3_ metabolism. To test these hypotheses in combination, we analysed intramolecular D abundances in leaf starch of the angiosperm *Helianthus annuus* synthesised at varying *C*_i_. In addition, we discuss the anaplerotic flux in the context of plant physiology and climate change.

## Material and Methods

### Samples

We used samples of the C_3_ plant *Helianthus annuus* L. cv. Zebulon from a previous study (Ehlers *et al.*, 2015). These plants were raised in 1.4 L pots in a greenhouse at an ambient CO_2_ concentration of *C*_a_≈450 ppm and a light intensity of 300-400 μmol photons m^−2^ s^−1^ (16 h photoperiod). They were watered daily, fertilised twice a week with Rika-S (Weibulls, Hammenhög, Sweden) and transferred to a growth chamber in groups of 8 after 7 to 8 weeks. Growth chamber temperature and relative humidity was set to 22/18 °C and 60/70% (day/night), respectively. On the first day after transfer, the plants were kept in darkness at *C*_a_=450 ppm to drain the starch reserves. During the following two days, the plant groups were grown at a *C*_a_ of either 180, 280, 450, 700, or 1500 ppm (300-400 μmol photons m^−2^ s^−1^, 16 h photoperiod). On the second day, gas exchange measurements were performed (Supporting Information, Notes S1), and leaves were harvested and stored at −20 °C until starch extraction described in Ehlers *et al.* (2015). Reported *C*_i_ values were estimated as described in the Supporting Information, Notes S2.

### Determination and expression of intramolecular deuterium abundances

Intramolecular D abundances in starch glucose were measured following published procedures (Betson *et al.*, 2006; Ehlers *et al.*, 2015). ^1^H NMR spectroscopy was used to ensure sample purity of the glucose derivative used for quantification (3,6-anhydro-1,2-*O*-isopropylidene-α-D-glucofuranose) of ≥99.5%. We recorded quantitative D NMR spectra on a DRX600 spectrometer with 5-mm broadband observe probe and ^19^F lock (Bruker BioSpin GmbH, Rheinstetten, Germany). Relative intramolecular D abundances were determined by signal deconvolution using Lorentzian line shape fits in TopSpin 3.1 (Bruker BioSpin GmbH, Rheinstetten, Germany). To screen for intramolecular fractionations, we expressed the data as deviations from the molecular average as

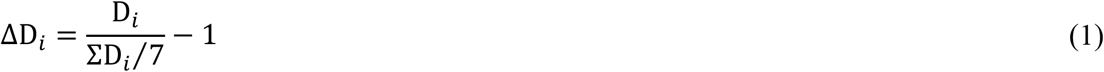

where D_*i*_ denotes relative D abundances at specific glucose hydrogen positions. In this notation, whole-molecule effects are not expressed. However, the size of any intramolecular fractionation effect is attenuated by its contribution to the denominator. To obtain estimates of effect sizes, we used as reference the average D abundance of the methyl-group hydrogens of the glucose derivative (introduced during derivatisation from a common batch of acetone) as

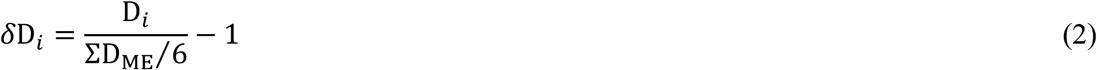

where D_ME_ denotes relative D abundances of the methyl-group hydrogens.

## Results

### Deuterium enrichments at glucose H^1^, and H^2^

Here, we analysed intramolecular D abundances in *Helianthus annuus* leaf starch synthesised at different levels of *C*_a_ (180, 280, 450, 700, 1500 ppm). We found intramolecular fractionations at H^1^ and H^2^ (Fig. 2). With decreasing *C*_a_ from 450 to 180 ppm (*C*_i_ from 328 to 140 ppm), *δ*D increases at H^1^ (210‰), and H^2^ (311‰, Fig. 3). Note that actual offsets may be somewhat larger than apparent offsets (Supporting Information, Notes S3). Above *C*_a_=450 ppm (*C*_i_≈328 ppm), these relationships break, i.e., they exhibit a change point at *C*_a_≈450 ppm. Generally, intramolecular fractionations are caused by metabolic processes (see below).

**Figure 2.**
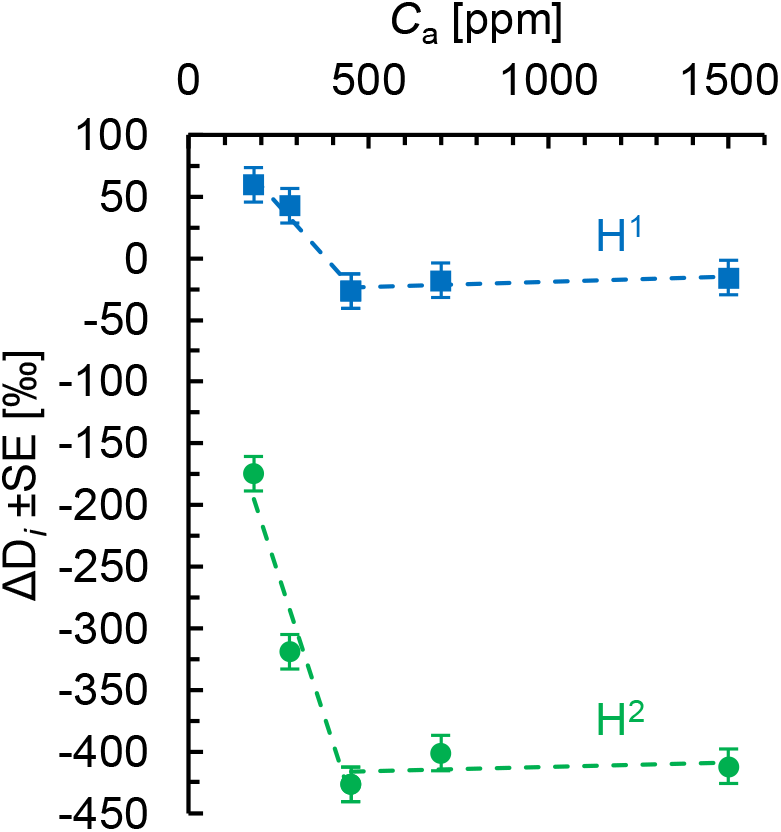
Relative D abundance at H^1^ (blue squares), and H^2^ (green circles) of *Helianthus annuus* leaf starch. Plants were grown in chambers at *C*_a_=450 ppm. After a day in darkness to drain starch reserves, plants were grown at different levels of *C*_a_ (180, 280, 450, 700, 1500 ppm) corresponding to different levels of *C*_i_ (140, 206, 328, 531, 1365 ppm) for two days. Data expressed in terms of the molecular average as ΔD_*i*_=D_*i*_/(ΣD_*i*_/7)-1 where D_*i*_ denotes relative D abundances at specific glucose hydrogen positions (±*SE*=14.1‰). Figure shows discrete data. Dashed lines added to guide the eye.

**Figure 3.**
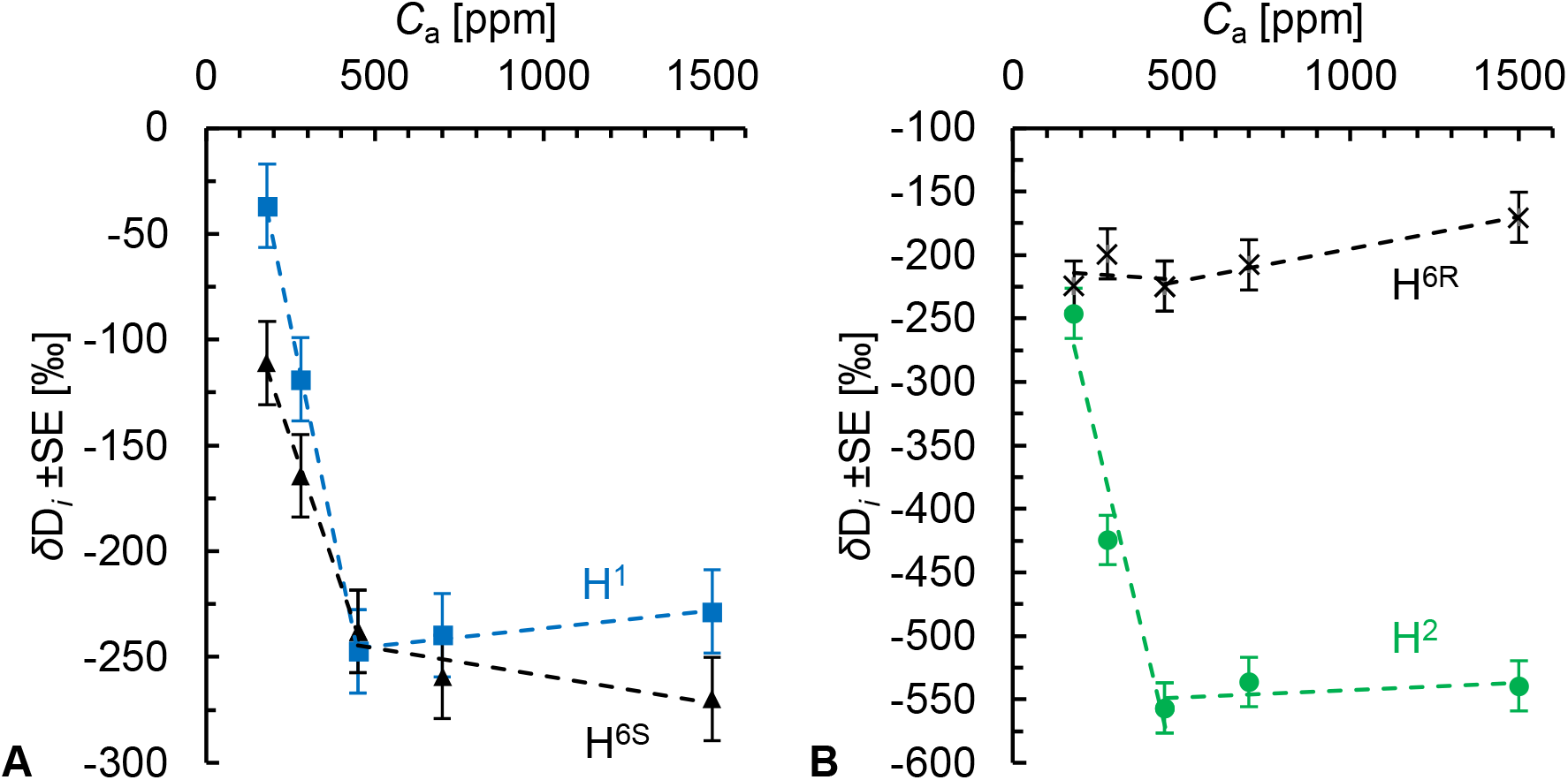
A-B Relative D abundance at H^1^ (blue squares), H^6S^ (black triangles), H^2^ (green circles), and H^6R^ (black crosses) of *Helianthus annuus* leaf starch. Plants were grown in chambers at *C*_a_=450 ppm. After a day in darkness to drain starch reserves, plants were grown at different levels of *C*_a_ (180, 280, 450, 700, 1500 ppm) corresponding to different levels of *C*i (140, 206, 328, 531, 1365 ppm) for two days. Data expressed as *δ*D_*i*_=D_*i*_/(ΣD_ME_/6)-1 with D_*i*_ and D_ME_ denoting relative D abundances at specific glucose hydrogen positions and the methyl-group hydrogens of the glucose derivative used for measurements, respectively (±*SE*=19.7‰). Figure shows discrete data. Dashed lines added to guide the eye.

### Photorespiratory deuterium fractionations occur at glucose H^1^, and H^6S^

With decreasing *C*_a_ from 450 to 180 ppm (*C*_i_ from 328 to 140 ppm), *δ*D increases at H^1^ (210‰), and H^6S^ (127‰) but not H^6R^ (Fig. 3). Changing photorespiration-to-photosynthesis ratios reportedly have equal effects on both the H^6R^/H^1^ and H^6R^/H^6S^ D abundance ratios (Schleucher, 1998). This was attributed to changes at H^6R^ (Schleucher, 1998). However, *δ*D_6R_ is invariable between 450 and 180 ppm (Fig. 3B). Thus, photorespiratory fractionation likely occurs at glucose H^1^ and H^6S^ but not H^6R^.

### Metabolic origin of the deuterium enrichment at glucose H^1^

Starch biosynthesis from photosynthetic glyceraldehyde 3-phosphate, GAP, proceeds via a multi-step enzymatic pathway (Fig. 4). In the first step, triosephosphate isomerase converts GAP (glucose C-4 to C-6) to dihydroxyacetone phosphate, DHAP (glucose C-1 to C-3). Thereby, the GAP hydrogen corresponding to glucose H^6S^ becomes the DHAP hydrogen corresponding to glucose H^1^. These hydrogens are neither modified by the triosephosphate isomerase reaction, nor by any subsequent reaction leading to starch. Hence, starch glucose H^6S^ and H^1^ can be expected to have equal D abundances. Evidently, this situation occurs at *C*_a_=450 ppm (Fig. 3A). Thus, there is no evidence for anaplerotic fractionation and flux at *C*_a_=450 ppm (under standard conditions) suggesting that G6PD is efficiently inhibited by thioredoxin despite the relatively low light intensity.

**Figure 4.**
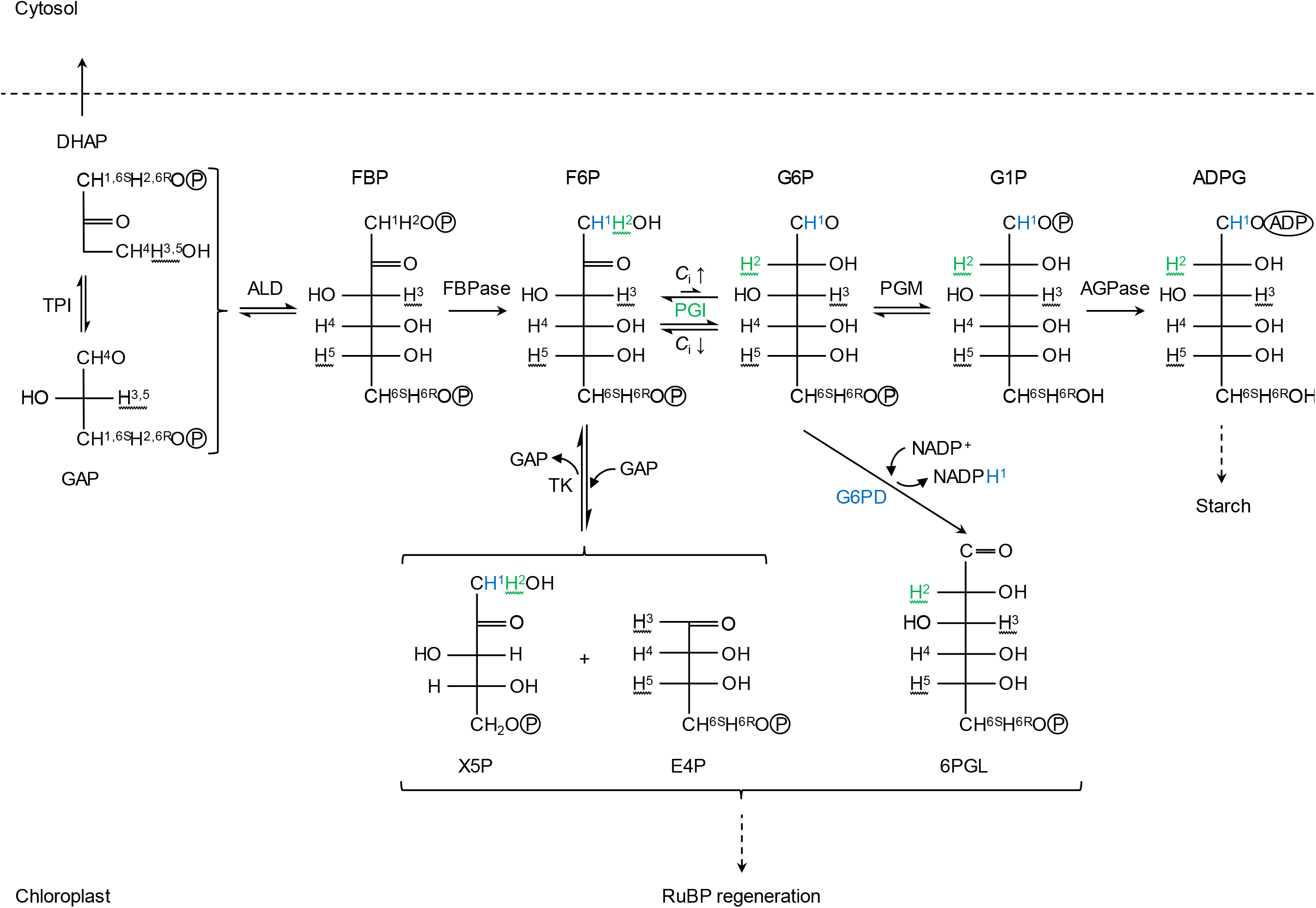
Hydrogen metabolism associated with starch biosynthesis. Hydrogens named according to their location(s) in starch glucose (indirect precursors not designated, see X5P). Green and blue hydrogens may be affected by isotope fractionation at PGI and G6PD, respectively. Wavy lines: Hydrogen from water is fractionally introduced during the TPI and PGI reaction. Under standard *C*_i_ conditions (*C*_i_↑), the chloroplastic PGI reaction is strongly displaced from equilibrium on the side of F6P. Under low *C*_i_ conditions (*C*_i_↓), the chloroplastic PGI reaction is closer to or at equilibrium. Dashed arrows: Intermediate reactions not shown. Abbreviations: 6PGL, 6-phosphogluconolactone; ADPG, ADP-glucose; AGPase, glucose-1-phosphate adenylyltransferase; ALD, fructose-1,6-bisphosphate aldolase; *C*_i_, intercellular CO_2_ concentration; DHAP, dihydroxyacetone phosphate; E4P, erythrose 4-phosphate; F6P, fructose 6-phosphate; FBP, fructose 1,6-bisphosphate; FBPase, fructose 1,6-bisphosphatase; G1P, glucose 1-phosphate; G6P, glucose 6-phosphate; G6PD, glucose-6-phosphate dehydrogenase; GAP, glyceraldehyde 3-phosphate; PGI, phosphoglucose isomerase; PGM, phosphoglucomutase; RuBP, ribulose 1,5-bisphosphate; TK, transketolase; TPI, triose-phosphate isomerase; X5P, xylulose 5-phosphate.

Equal D enrichments at H^1^ and H^6S^ can be attributed to photorespiratory fractionation (see above). However, at *C*_a_=180 ppm, H^1^ is 74‰ more enriched than H^6S^ (Fig. 3A; *δ*D_1_=−37‰, *δ*D_6S_=−111‰). This enrichment can be expected to be introduced beyond DHAP synthesis by a high-flux carrying pathway branching off from the starch biosynthesis pathway. To our knowledge, there are only two possible candidates, the well-established CBC-regeneration pathway and the proposed anaplerotic pathway (Fig. 4). The former pathway starts with the conversion of F6P and GAP to erythrose 4-phosphate and xylulose 5-phosphate by transketolase. The hydrogen corresponding to starch glucose H^1^ is not modified by the transketolase reaction. By contrast, the anaplerotic pathway starts with the irreversible G6PD-catalysed conversion of G6P to 6-phosphogluconolactone which deprotonates the G6P hydrogen corresponding to starch glucose H^1^. This reaction reportedly has a deuterium isotope effect *in vitro* (*α*_D_=*k*_H_/*k*_D_=2.97) (Hermes *et al.*, 1982). Thus, G6PD is a plausible candidate for D fractionation at H^1^.

Consistent with our observations, fractionation modelling shows that increasing anaplerotic flux, *f*, causes D increases at H^1^ of remaining G6P, *δ*D_G6P’_ (Fig. 5; Supporting Information, Notes S4). A 74‰ increase as observed in starch (Fig. 3A) indicates that ≈10% of the G6P entering the starch biosynthesis pathway is diverted into the anaplerotic pathway. However, G6P can be converted back to F6P by PGI and enter the CBC via transketolase (Fig. 4). This flux will be low under low chloroplastic G6P concentrations (i.e., under standard growth conditions) yet increase as the PGI reaction shifts towards equilibrium (with shifts towards low *C*_a_, see ‘Introduction’). It can be expected to fractionally remove the G6PD-fractionation signal from the starch biosynthesis pathway. Thus, at *C*_a_=180 ppm, more than 10% of the G6P entering the starch biosynthesis pathway may be diverted into the anaplerotic pathway (to obtain a 74‰ increase at H^1^).

**Figure 5.**
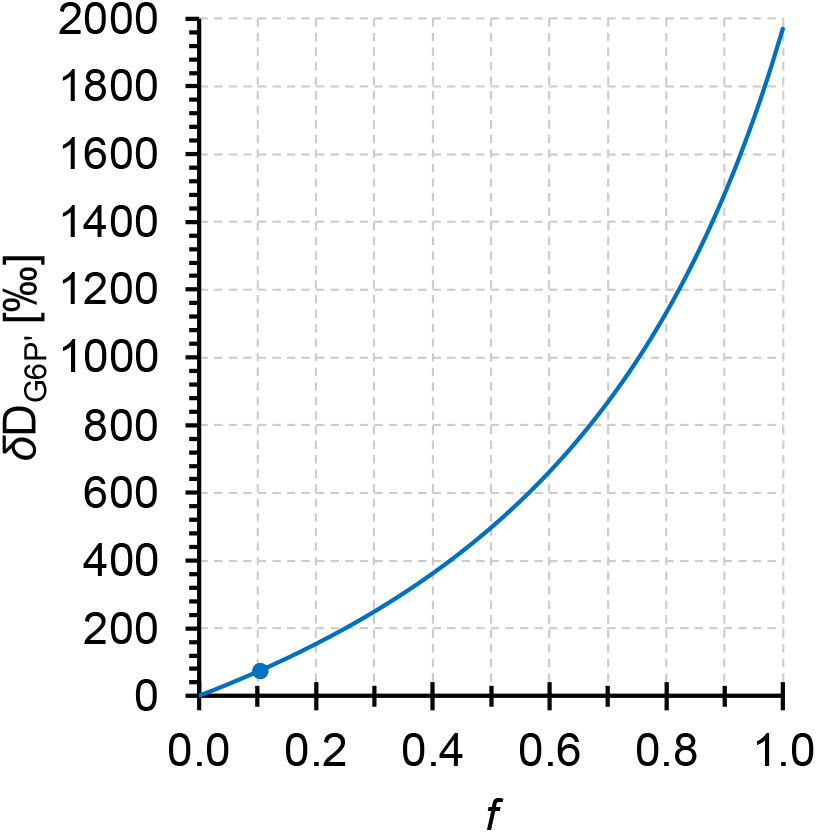
Modelled D fractionation by glucose-6-phosphate dehydrogenase (G6PD) at H^1^ of glucose 6-phosphate, *δ*D_G6P’_. Modelling assumed an open system at steady state. Incoming G6P has two fates, starch biosynthesis or anaplerotic reinjection into the CBC via G6PD (Fig. 4). Commitment of G6P to the anaplerotic pathway vs. starch biosynthesis, *f*, is given on the abscissa (e.g., *f*=1: all G6P enters the anaplerotic pathway, *f*=0: all G6P enters starch biosynthesis). Blue dot: *f*=0.1044, *δ*D_G6P’_=74.41‰. G6PD has an *in vitro* kinetic isotope effect of *α*_D_=2.97 (Hermes *et al.*, 1982). G6P entering the system was assumed to have the same D/H ratio as Vienna Standard Mean Ocean Water (*R*_VSMOW_=;155.76*10^−6^) (Hagemann *et al.*, 1970). Data expressed in terms of *R*_VSMOW_.

### Metabolic origin of the deuterium enrichment at glucose H^2^

At the level of chloroplastic triose phosphates, the precursors of starch glucose H^2^ correspond to the precursors of starch glucose H^6R^ (Fig. 4). In the biosynthetic pathway from triose phosphate to starch, the former is subject to modification by PGI while the latter is not modified. Therefore, we assume the D abundance at H^6R^ is not modified during starch biosynthesis and can be used as reference for H^2^.

Under standard conditions (*C*_a_=450 ppm), starch glucose H^2^ is 311‰ depleted compared to low *C*a conditions (=180 ppm) and 332‰ depleted compared to H^6R^ (Fig. 3B). This is consistent with reported glucose H^2^ depletions in leaf starch compared to sucrose (*Phaseolus vulgaris*, 333‰; *Spinacia oleracea*, 500‰) which were explained by different *modus operandi* of chloroplastic and cytosolic PGI (Schleucher *et al.*, 1999). Reportedly, the chloroplastic PGI reaction is strongly displaced from equilibrium on the side of F6P (Dietz, 1985; Gerhardt *et al.*, 1987; Leidreiter *et al.*, 1995; Schleucher *et al.*, 1999) while the cytosolic reaction is either somewhat displaced from equilibrium (Leidreiter *et al.*, 1995; Schleucher *et al.*, 1999) or at equilibrium (Gerhardt *et al.*, 1987). Thus, in chloroplasts under standard conditions, the kinetic isotope effect of PGI is manifested. This effect can cause D depletions of up to ≈550‰ (Supporting Information, Notes S5; hydrogen exchange with the medium not considered).

At low *C*_a_ (=180 ppm), starch glucose H^2^ is merely 21.5‰ depleted compared to H^6R^ (Fig. 3B). This is consistent with a shift of the PGI reaction from kinetic to equilibrium conditions and thus upregulated anaplerotic flux (see ‘Introduction’). Theoretically, equilibrium isotope fractionation by PGI causes a D depletion of ≈99‰ when fractionation related to hydrogen exchange with the medium is not considered (Supporting Information, Notes S5). However, the offset between theoretical and observed values (≈77.5‰) indicates that hydrogen exchange with the medium affects isotope fractionation by PGI.

## Discussion

Consistent with isotope theory related to anaplerotic carbon flux into the CBC (see ‘Introduction’; Wieloch *et al.*, 2021), we found D fractionation signals at H^1^ and H^2^ of leaf starch which increase with decreasing *C*_a_ (and *C*_i_) below a response change point (Figs. 2–3). Under standard conditions (*C*_a_=450 ppm, *C*_i_=328 ppm), we estimate negligible anaplerotic flux. This is in contrast to findings by Xu *et al.* (2021). At *C*_a_=180 ppm (*C*_i_=140 ppm), we estimate that more than 10% of the G6P entering the starch biosynthesis pathway is diverted into the anaplerotic pathway. Previously, we reported evidence for anaplerotic flux in *Pinus nigra*, a gymnosperm (Wieloch *et al.*, 2018, 2021). Here, we report evidence for this flux in the angiosperm *Helianthus annuus* (Figs. 2–3). Thus, anaplerotic flux into the CBC may be a general component of C3 metabolism. Lastly, our data suggest photorespiratory fractionation affects the D abundance at glucose H^1^ and H^6S^ but not H^6R^.

### Anaplerotic flux into the CBC: A leaf-level response to source-sink imbalances?

In *Helianthus annuus*, anaplerotic flux was upregulated under apparent steady-state conditions at low *C*_a_ concentrations (Figs. 2–3). Similarly, large anaplerotic isotope fractionations in *Pinus nigra* tree rings (formed over the course of several months) suggest that the anaplerotic flux can persist over long periods (Wieloch *et al.*, 2018, 2021).

*Helianthus annuus* was raised at *C*_a_=450 ppm over 7 to 8 weeks. After a day in darkness, the plants were subjected to different *C*_a_ treatments for two days. We found evidence for anaplerotic flux at *C*_a_<450 ppm (Figs. 2–3). This may be explained by a shift from the plants’ previously established source-sink balance at *C*_a_=450 ppm to source-limited conditions at *C*_a_<450 ppm (where sink demands exceed the source strength). A concomitant downstream pull of carbohydrates may have caused excess export of triose phosphates from chloroplasts to the cytosol and shortage of chloroplastic triose phosphates for regeneration of ribulose 1,5-bisphosphate, RuBP (Fig. 4). To maintain the photosynthetic/photorespiratory capacity at the level of RuBP, anaplerotic flux may then have been upregulated by mechanisms described above (see ‘Introduction’). Thus, we propose upregulation of the anaplerotic flux is a leaf-level response to source-limited growth conditions. It may be triggered by all environmental parameters that can cause source limitations such as low *C*_a_, low *C*_i_, and drought (as in *Pinus nigra*) (Wieloch *et al.*, 2018, 2021). This may have implications for gas exchange studies aiming to investigate actual steady-state conditions. To preclude errors due to anaplerotic CO_2_ liberation associated with disturbed source-sink balances, plants should be raised under the same conditions they are analysed.

### Anaplerotic flux into the CBC: Potential benefits for plant functioning and growth

Like photorespiration, anaplerotic flux has a negative carbon balance (Fig. 1) and seems, therefore, detrimental for plant functioning and growth but we propose two potential benefits.

First, by maintaining RuBP levels, anaplerotic flux maintains photorespiration (both fluxes increase with decreasing *C*_a_ and *C*_i_). Reportedly, photorespiration is positively correlated with *de novo* nitrate assimilation into protein (Bloom, 2015). Thus, anaplerotic flux may help to increase photosynthetic capacity at the enzyme level, and it seems plausible that plants would try to increase this capacity under source-limited conditions. Additionally, increased nitrogen assimilation may support the biosynthesis of nitrogen-containing compounds used for maintenance and repair processes. Overall, it seems plausible that metabolism would shift towards nitrogen assimilation when carbon metabolism is substrate limited.

Second, optimal plant functioning requires a balance between energy supply and consumption (Huner *et al.*, 1996). Upregulation of the anaplerotic flux occurs at low *C*_i_ (Figs. 2–3) which can cause energy imbalances in chloroplasts (Huner *et al.*, 1996; Wilhelm & Selmar, 2011; Vanlerberghe *et al.*, 2015). This involves decreased consumption of ATP and NADPH by the CBC but constant electron input by light-harvesting complexes. The resulting lack of electron acceptors (NADP^+^ and ADP) leads to the generation of reactive oxygen species (Vanlerberghe *et al.*, 2015). Futile carbon cycling involving CO_2_ liberation by the anaplerotic flux and refixation by Rubisco has a negative NADPH and especially ATP balance (Fig. 1). Thus, anaplerotic flux may help to dissipate excess energy and counteract oxidative stress and photoinhibition at low *C*_i_.

In *Pinus nigra*, upregulation of the anaplerotic pathway was reportedly associated with below-average yet not exceptionally low growth (Wieloch *et al.*, 2021). This is surprising since anaplerotic flux increases with decreasing *C*_i_ and liberates CO_2_ (Figs. 1–3). Principally, inputs of storage carbohydrates from previous years may have rescued growth rates but such inputs were shown to be negligible (Wieloch *et al.*, 2018). By contrast, physiological benefits proposed above may have contributed to maintaining growth rates.

### Anaplerotic flux into the CBC and climate change

In *Pinus nigra*, anaplerotic flux reportedly correlates positively with drought and negatively with *C*_a_ (Wieloch *et al.*, 2018, 2021). This is consistent with findings reported here. The Intergovernmental Panel on Climate Change predicts a higher frequency of drought events in already dry regions towards the end of the 21^st^ century (RCP8.5) (IPCC, 2021). Simultaneously, *C*_a_ will increase globally. Therefore, it is unclear whether photosynthesis involving an upregulated anaplerotic flux will become more prevalent.

Despite potentially significant impacts on carbon, nitrogen, and energy metabolism, the anaplerotic pathway is currently neither well understood nor considered in models of photosynthesis, biosphere-atmosphere CO_2_ exchange, and plant growth. Stable isotope tools utilised here may enable comprehensive flux analyses across spatiotemporal scales.

## Author contributions

T.W. conceived the study and led the research. A.A., and J.S. prepared the samples and acquired the data. T.W. analysed and interpreted the data. T.W. wrote the paper with input from A.A., and J.S.

## Data availability

The data that support the findings of this study are available from the corresponding author upon reasonable request.

