## Supporting information for "Anaplerotic flux into the Calvin-Benson cycle. Hydrogen isotope evidence for *in vivo* occurrence in C_3_ metabolism"

### Notes S1. Leaf gas exchange measurements

Gas exchange measurements were performed with a two-channel fast-response system described previously (Laisk & Edwards, 1997). A sandwich-type round chamber enclosed a leaf area of 8.04 $cm^2$. Gas flow was maintained at 0.5 mmol $s^{-1}$. Measurements were performed at $CO_2$ concentrations similar to those applied in the growth chamber over two days ($C_a$=173 ± 0.1SE, 269 ± 0.1SE, 433 ± 0.1SE, 675 ± 0.2SE, 1431 ± 0.4SE ppm; n=8). To maintain temperatures of 22 ± 0.5 °C, the upper leaf side was attached to a thermostated glass window by starch gel. Light was provided by a Schott KL 1500 lamp (Walz Heinz GmbH, Effeltrich, Germany) and measured by a LI-185 quantum sensor (LiCor, Lincoln, USA). Measurements were performed at 300 µmol photons $m^{-2}$ $s^{-1}$ yielding $C_i$=136 ± 1SE, 202 ± 4SE, 304 ± 8SE, 516 ± 14SE, 1282 ± 16SE ppm.

### Notes S2. Estimation of intercellular $CO_2$ concentrations during growth chamber experiments

Figure S1 shows $C_a$ and $C_i$ data obtained by gas exchange analysis (black circles). Their relationship is well described by a quadratic equation (dotted line, $R^2$=0.999, $p$<0.001, $n$=5). Based on this equation, we estimated $C_i$ values of 140, 206, 328, 531, and 1365 ppm for growth chamber treatments of $C_a$=180, 280, 450, 700, and 1500 ppm (red circles).

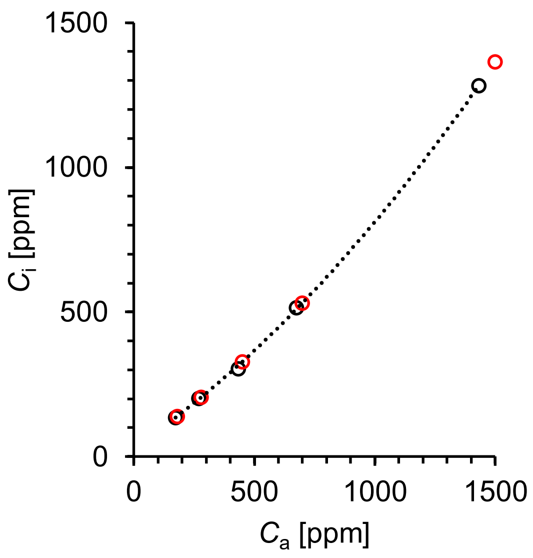


**Figure S1** Ambient and intercellular $CO_2$ concentrations in *Helianthus annuus* leaves ($C_a$, and $C_i$, respectively). Black circles: measured values obtained by gas exchange analysis. Black dotted line: quadratic relationship between $C_a$ and $C_i$ measurements ($R^2$=0.999, $p$<0.001, $n$=5). Red circles: $C_i$ values during growth chamber experiments estimated based on the quadratic equation.

**Notes S3. Dilution of isotope signals by remnant starch**

The plants studied here were grown at a $CO_2$ concentration of 450 ppm over 7 to 8 weeks. $H^1$ and $H^2$ of starch glucose synthesised under these conditions have $\delta$D values of -247‰ and -557‰, respectively (Fig. 3). To drain the starch reserves and avoid dilution by 450 ppm isotope signals, all plants were kept in darkness for 24 hours after transfer to growth chambers. Reportedly, this treatment led to a starch reduction from about 8 to almost 0 μmol glucose $g^{-1}$ FW in *Arabidopsis thaliana* (Smith & Stitt, 2007). Similarly, we found a starch remnant of <0.6 μmol glucose $g^{-1}$ FW in our samples. After two days of acclimation to 1500 ppm, we found a starch content of >171 μmol glucose $g^{-1}$ FW. Thus, at 1500 ppm $CO_2$, dilution of isotope signals due to remnant (450 ppm) starch is negligible (0.6/171=4‰).

At 180 ppm, net $CO_2$ assimilation was lower than at 1500 ppm (5.1 vs. 17.3 μmol $m^{-2}$ $s^{-1}$). Assuming equal relative carbon partitioning under both conditions, dilution of isotope signals in starch synthesised at 180 ppm is small (0.6/(171*5.1/17.3)=1.2%). $H^1$ and $H^2$ of starch glucose synthesised at 180 ppm have apparent $\delta$D values of -37‰ and -246‰, respectively (Fig. 3). Accounting for a 1.2% dilution by remnant starch, we estimate actual $\delta$D values of -34.4‰ and -242.2‰, respectively. However, sucrose-to-starch carbon partitioning ratios increase as ambient $CO_2$ concentrations decrease (Sharkey et al. 1985). Thus, at 180 ppm $CO_2$, the relative contribution of remnant (450 ppm) starch may be larger, i.e., actual $\delta$D values and anaplerotic flux may be somewhat higher than estimated.

**Notes S4. Deuterium fractionation by glucose-6-phosphate dehydrogenase**

Modelling of fractionations by glucose-6-phosphate dehydrogenase, G6PD, assumed an open system at steady state and followed published basic procedures (Hayes, 2002). Incoming glucose 6-phosphate, G6P, has two fates, starch biosynthesis or anaplerotic reinjection into the Calvin-Benson cycle via G6PD. D fractionation between the reaction product, 6-phosphogluconolactone (6PGL), and remaining educt, G6P', is given as

$$R_{\mathrm{6PGL}} = \frac{1}{\alpha} R_{\mathrm{G6P\prime}} \tag{S1}$$

where $R$ denotes D/H ratios, and $\alpha$ denotes D isotope effects of G6PD ($\alpha_D$=2.97) (Hermes *et al.*, 1982). Isotope mass balance of the system is given as

$$F_{\mathrm{G6P}} = fF_{\mathrm{6PGL}} + (1-f)F_{\mathrm{G6P'}} \quad \text{(S2)}$$

where $F$ denotes fractional abundances (D/(H+D)), and $f$ denotes the 6PGL commitment ($f$ was varied between 0 and 1). $F$ and $R$ relate to each other as

$$F = \frac{R}{1+R} \quad \text{(S3)}$$

Substituting $F$ for $R$ in equation S2 yields

$$\frac{R_{\mathrm{G6P}}}{1+R_{\mathrm{G6P}}} = f\frac{R_{\mathrm{6PGL}}}{1+R_{\mathrm{6PGL}}} + (1-f)\frac{R_{\mathrm{G6P'}}}{1+R_{\mathrm{G6P'}}} \quad \text{(S4)}$$

Here, we substitute the unknown D/H ratio of incoming G6P, $R_{\mathrm{G6P}}$, by the known ratio of Vienna Standard Mean Ocean Water, $R_{\mathrm{VSMOW}}$ (155.76*$10^{-6}$) (Hagemann *et al.*, 1970). Consequently, fractionations expressed on the $\delta$-scale (Eq. S7) develop from zero. Substituting $R_{\mathrm{6PGL}}$ in equation S4 based on equation S1 then gives

$$\frac{R_{\mathrm{VSMOW}}}{1+R_{\mathrm{VSMOW}}} = f\frac{R_{\mathrm{G6P'}}/\alpha}{1+R_{\mathrm{G6P'}}/\alpha} + (1-f)\frac{R_{\mathrm{G6P'}}}{1+R_{\mathrm{G6P'}}} \quad \text{(S5)}$$

Solving equation S5 for $R_{\mathrm{G6P'}}$ gives

$$c = \frac{-ad - a + b + d - bd \pm Z}{2(a-1)} \quad \text{(S6)}$$

with

$$Z = \sqrt{a^2d^2 - 2ad^2 + b^2d^2 + 2abd^2 - 2bd^2 + d^2 - 2a^2d + 2ad - 2b^2d + 2bd + a^2 + b^2 - 2ab}$$

where $a$ to $d$ denote $R_{\mathrm{VSMOW}}/(1+R_{\mathrm{VSMOW}})$, $f$, $R_{\mathrm{G6P'}}$, and $\alpha$, respectively. While there are two mathematically correct solutions (±$Z$), only one of them returns plausible fractionation values (-$Z$). Modelling results were expressed in terms of $R_{\mathrm{VSMOW}}$ as

$$\delta \mathrm{D}_{\mathrm{G6P'}} = \frac{R_{\mathrm{G6P'}}}{R_{\mathrm{VSMOW}}} - 1 \qquad \text{(S7)}$$

**Notes S5. Deuterium fractionation by phosphoglucose isomerase**

In the F6P to G6P direction, spinach PGI has a tritium isotope effects, $\alpha_{\mathrm{T}}$, of 3 (Noltmann, 1972). The corresponding deuterium isotope effect, $\alpha_{\mathrm{D}}$, can be estimated from $\alpha_{\mathrm{T}}$ as

$$\alpha_{\mathrm{D}} = k_{\mathrm{H}}/k_{\mathrm{D}} = \sqrt[1.44]{\alpha_{\mathrm{T}}} \qquad \text{(S8)}$$

where $k_{\mathrm{H}}$ and $k_{\mathrm{D}}$ denote the reaction rates of the protium and deuterium isotopologues of F6P, respectively (Melander & Saunders, 1980). Based on this relationship, we estimate an $\alpha_{\mathrm{D}}$ of 2.14 which may cause D depletions at G6P $H^2$ of about 534‰ ($1/\alpha_{\mathrm{D}}$-1).

For rabbit muscle PGI, Rose and O'Connell (1961) measured an $\alpha_{\mathrm{D}}$ of 2.22 in the F6P to G6P direction (Table S1) which is similar to the $\alpha_{\mathrm{D}}$ estimated for spinach PGI. In the G6P to F6P direction, these authors measured an $\alpha_{\mathrm{D}}$ of 2. From these values, we calculated an equilibrium isotope effect, $EIE_{\mathrm{D}}$, of 1.11 in G6P. The kinetic isotope of effect (F6P to G6P direction, $\alpha_{\mathrm{D}}$=2.22) may cause D depletions at G6P $H^2$ of about 550‰ while the equilibrium isotope effect ($EIE_{\mathrm{D}}$=1.11) may cause D depletions of about 99‰ ($1/EIE_{\mathrm{D}}$-1). Hence, with a shift from kinetic to equilibrium conditions, the D abundance at G6P $H^2$ may increase by about 451‰.

**Table S1** *In vitro* D isotope effects of PGI (Rose & O'Connell, 1961).

| F6P | | G6P |
|---|---|---|
| | → | 2.22 |
| 0.9 | ↔ | 1.11 |
| 2 | ← | |

→ kinetic isotope effect, ↔ equilibrium isotope effect. Calculated values underlined.

Note, our discussion implicitly assumes that all hydrogen from F6P $H^{1R}$ is transferred to G6P $H^2$. However, at 22 °C, spinach PGI transfers only 70% of the hydrogen while the rest undergoes exchange with the medium (F6P to G6P direction) (Noltmann, 1972). To our knowledge, isotope effects of hydrogen loss to the medium and hydrogen uptake from the medium are unknow as is the hydrogen isotope composition of the medium. Furthermore, hydrogen exchange rates are temperature dependent and differ between the forward and backward reaction of PGI (Rose & O'Connell, 1961; Noltmann, 1972). Thus, isotope fractionation by hydrogen exchange requires further attention.
